# Upregulation of the Cdc42 GTPase limits the replicative lifespan of budding yeast

**DOI:** 10.1101/2021.04.27.441634

**Authors:** Pil Jung Kang, Rachel Mullner, Haoyu Li, Derek Hansford, Han-Wei Shen, Hay-Oak Park

## Abstract

Cdc42, a conserved Rho GTPase, plays a central role in polarity establishment in yeast and animals. Cell polarity is critical for asymmetric cell division, and asymmetric cell division underlies replicative aging of budding yeast. Yet how Cdc42 and other polarity factors impact lifespan is largely unknown. Here, we show by live-cell imaging that the active Cdc42 level is sporadically elevated in wild type during repeated cell divisions but rarely in the long-lived *bud8* deletion cells. We find a novel Bud8 localization with cytokinesis remnants, which also recruit Rga1, a Cdc42 GTPase activating protein. Genetic analyses and live-cell imaging suggest that Rga1 and Bud8 oppositely impact lifespan likely by modulating active Cdc42 levels. An *rga1* mutant, which has a shorter lifespan, dies at the unbudded state with a defect in polarity establishment. Remarkably, Cdc42 accumulates in old cells, and its mild overexpression accelerates aging with frequent symmetric cell divisions, despite no harmful effects on young cells. Our findings implicate that the interplay among these positive and negative polarity factors limits the lifespan of budding yeast.

## INTRODUCTION

While budding yeast cells undergo a finite number of cell divisions, the yeast population is immortal in the presence of nutrients due to asymmetric cell division. Replicative lifespan (**RLS**) is defined by the number of daughter cells a mother cell produces before senescence or death (Steinkraus *et al*., 2008).

Each cell division leaves a chitin-rich ‘bud scar’ and the chitin-less ‘birth scar’ on the mother and daughter cell surface, respectively (Barton, 1950; Mortimer and Johnston, 1959). These remnants of cell division (i.e., ‘cytokinesis remnants’, **CRM**s) are associated with two interdependent transmembrane proteins Rax1 and Rax2 (Chen *et al*., 2000; Kang *et al*., 2004) (Fig. 1A). Since yeast cells bud at a new site that does not overlap with any previous bud site, the number of bud scars reflects the replicative age. Earlier reports postulated that depletion of available cell surface or nutrient exchange with the environment because of bud scars might limit RLS (Mortimer and Johnston, 1959; Johnston, 1966). Later studies, however, disapproved of this idea because aging does not correlate with the amount of chitin on the cell wall. Furthermore, daughter cells born from old mothers often have reduced lifespan despite the absence of bud scars (Egilmez and Jazwinski, 1989; Kennedy *et al*., 1994; Nestelbacher *et al*., 1999). Instead, aging appears to involve conserved regulatory systems across eukaryotic species (Kenyon, 2001; Wasko and Kaeberlein, 2014; McCormick *et al*., 2015).

**Figure 1.**
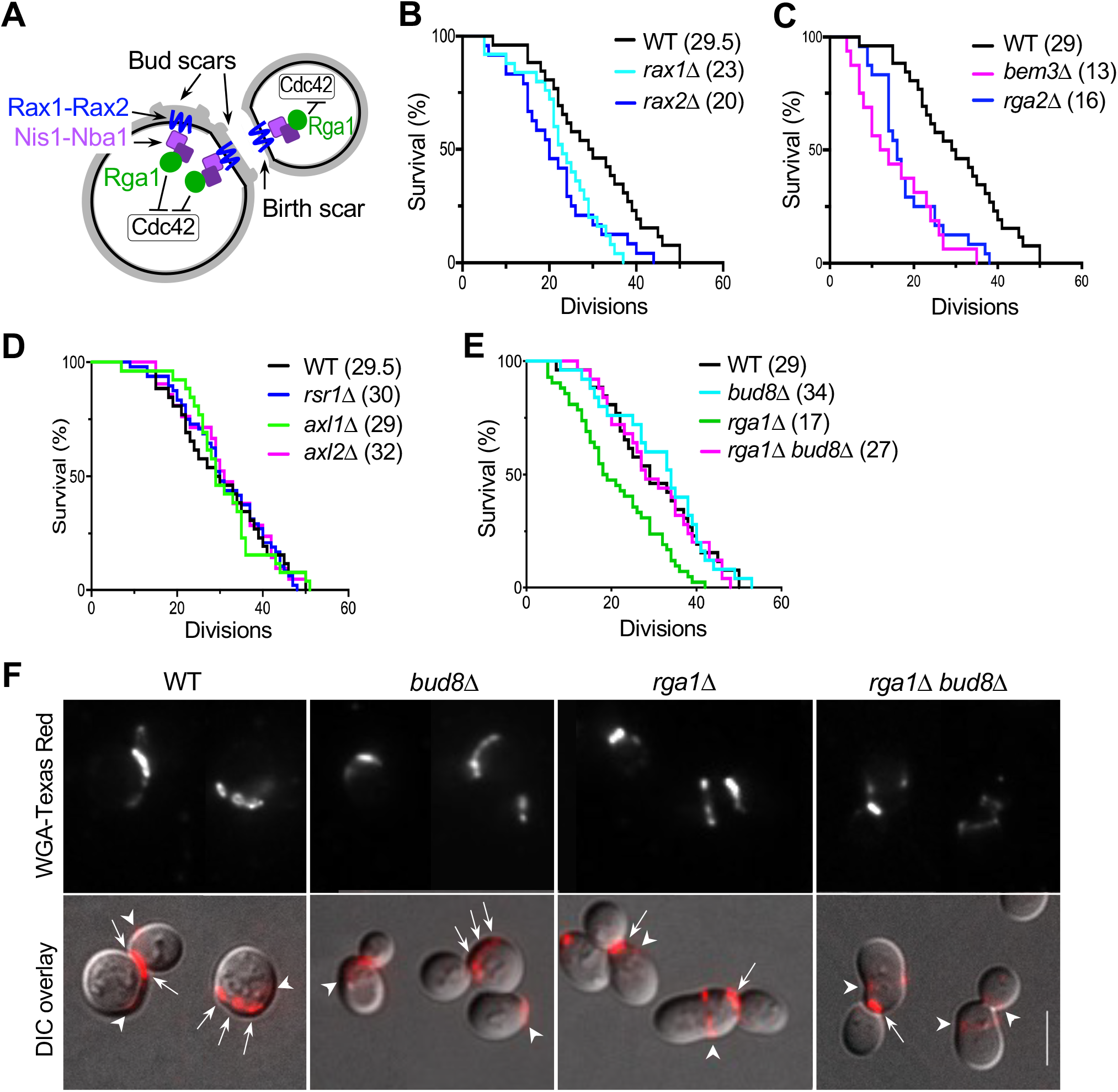
RLS measurement and characterization of a *bud8*Δ *rga1*Δ mutant. **A**. A scheme shows the negative polarity factors associated with the Rax1-Rax2 complex at CRMs (i.e., the birth scar in daughter cells and bud scars in mother cells). Rga1 at CRMs inhibits Cdc42 activation at any previous cell division site. **B – E**. Graphs show the percentage of surviving cells after the indicated number of divisions. All strains have the same background (see Table S1). The median survival age of each strain is indicated in parentheses. *p* values are obtained from Log-rank (Mantel-Cox) test. **B**. WT (*n* = 28), *rax1*Δ (*n* = 25), and *rax2*Δ (*n* = 25). *p* = 0.0039 (WT vs *rax1*Δ) and 0.0070 (WT vs *rax2*Δ). **C**. WT (*n* = 26), *bem3*Δ (*n* = 20), and *rga2*Δ (*n* = 24). p < 0.0001 (WT vs *bem3*Δ; and WT vs *rga2*Δ). **D**. WT (*n* = 28), *rsr1*Δ (*n* = 48), *axl1*Δ (*n* = 26), and *axl2*Δ (*n* = 31). *p* = 0.84 (WT vs *rsr1*Δ), 0.97 (WT vs *axl1*Δ), and 0.89 (WT vs *axl2*Δ). **E**. WT (*n* = 26), *rga1*Δ (*n* = 47), *bud8*Δ (*n* = 25), and *bud8*Δ *rga1*Δ (*n* = 27). *p* = 0.0004 (WT vs *rga1*Δ), 0.0026 (*rga1*Δ *bud8*Δ vs *rga1*Δ), 0.54 (WT vs *bud8*Δ *rga1*Δ), and 0.74 (WT vs *bud8*Δ). **F**. WGA-Texas Red staining of WT and each mutant (n = 80 cells for each strain). Bud scars (marked with arrows) are usually smaller and show stronger fluorescence than the birth scar (marked with arrowheads). Fluorescence images were deconvolved. Scale bar, 5 µm.

Yeast cells undergo oriented cell division in two distinct patterns depending on their cell type. Haploid **a** or α cells bud in the axial pattern, in which both mother and daughter cells select a new bud site adjacent to their immediately preceding division site. Diploid **a**/ α cells bud in the bipolar pattern, in which daughter cells bud preferentially at the distal pole (*i*.*e*., pole distal to the birth scar), and mother cells choose a bud site near either pole. The selection of a proper bud site requires Bud proteins, which recruit and activate the Cdc42 GTPase module and thus determine the orientation of the polarity axis for budding (Bi and Park, 2012; Kang *et al*., 2014). Rga1, a Cdc42 GTPase-activating protein (**GAP**), is involved in establishing the proper axis of cell polarity by preventing Cdc42 repolarization within the previous division site (Tong *et al*., 2007). Rga1 also localizes to CRMs by direct interaction with Nba1 and Nis1 (Miller *et al*., 2017), negative polarity cues that are recruited by Rax1-Rax2 (Meitinger *et al*., 2014) (Fig. 1A). Cells lacking the negative polarity factors, such as Rga1 and Nba1, have a shorter RLS than wild type (**WT**) (Meitinger *et al*., 2014).

Interestingly, a large-scale study of yeast deletion mutants identified *bud8*Δ as one of the long-lived mutants (McCormick *et al*., 2015). Bud8 is involved in bipolar budding but has no known function in **a** or α cells (Zahner *et al*., 1996). Despite these relatively well-known roles of the polarity proteins in polarized growth, whether and how these proteins play any causative role in aging remains elusive. To address these outstanding questions, we performed genetic analysis and single-cell imaging for the entire lifespan of budding yeast. Our findings suggest that upregulation of Cdc42 activity and level limits RLS and that Bud8 and Rga1 oppositely impact lifespan likely by modulating the active Cdc42 level.

## RESULTS & DISCUSSION

Deletion mutants of the negative polarity factors, *rga1*Δ and *nba1*Δ, often bud within the old cell division site, resulting in a narrow bud neck (Tong *et al*., 2007; Meitinger *et al*., 2014). This phenotype is thought to cause a shorter RLS through nuclear segregation defects (Meitinger *et al*., 2014). Since Rax1 and Rax2 recruit Rga1 via the Nba1-Nis1 complex (Miller *et al*., 2017), we asked whether *rax1*Δ and *rax2*Δ mutants also have shorter RLS. By a micromanipulation-based assay, we found that *rax1*Δ and *rax2*Δ mutants have shorter RLS than WT (Fig. 1B) despite little growth defect during the exponential growth phase. Unlike *rga1* cells, *rax1*Δ and *rax2*Δ cells often bud normally at a site adjacent to the previous division site, except the first budding event, which occurs within the birth scar (Miller *et al*., 2017), and thus these cells do not have the narrow bud neck. These observations raised the question of whether another defect of *rga1*Δ mutants might be a cause of a shorter RLS.

We wondered whether the lack of (or reduced) Cdc42 GAP activity of *rga1*Δ mutants might result in a shorter RLS. Indeed, when we examined other Cdc42 GAP mutants *rga2*Δ and *bem3*Δ, these mutants also displayed significantly reduced RLS than WT (Fig. 1C). The *rga2*Δ and *bem3*Δ cells do not bud within the previous division site and thus do not have a narrow bud neck (Tong *et al*., 2007). These findings led us to speculate that Cdc42 activity might impact RLS. We also considered another possibility that a shorter RLS of *rga1*Δ mutants might be due to its bud-site selection defect (Smith *et al*., 2002; Tong *et al*., 2007). Some physiological changes observed during yeast aging, such as budding pattern alteration, have been proposed to be a cause rather than a consequence of aging (Jazwinski *et al*., 1998). However, all three *bud* mutants tested – *rsr1*Δ, *axl1*Δ, and *axl2*Δ – had similar RLS as WT (Fig. 1D). Interestingly, however, a haploid *bud8*Δ mutant has a longer RLS than WT (Fig. 1E), as previously reported (McCormick *et al*., 2015). Since Bud8 is dispensable for the axial budding pattern of **a** or cells (Zahner *et al*., 1996), the longer RLS of *bud8*Δ mutants is likely to be independent of its role in bud-site selection. These results suggest that any alteration of the orientation of the polarity axis *per se* is unlikely to affect replicative aging.

Since deletions of *BUD8* and *RGA1* have opposite effects on RLS, we examined their genetic interactions. When we measured RLS of a *bud8*Δ *rga1*Δ double mutant and each single mutant, we found that the median RLS of the *bud8*Δ *rga1*Δ mutant was similar to that of WT (Fig. 1E). A *bud8*Δ *rga1*Δ mutant divided significantly more than *rga1*Δ but less than *bud8*Δ before death. We then asked whether deletion of *BUD8* also rescues the same-site re-budding phenotype of *rga1* mutants by staining the birth scar and bud scars with WGA (Wheat germ agglutinin)-Texas Red. Over 90% of WT and *bud8*Δ cells showed bud scars or buds next to the birth scar, as expected for cells budded in the axial pattern. In contrast, about 90% of *bud8*Δ *rga1*Δ cells had buds or bud scar(s) within the birth scar, similar to that of *rga1*Δ cells (Fig. 1F). These results indicate that deletion of *BUD8* rescues a shorter RLS, but not the same-site re-budding phenotype, of *rga1*Δ mutants. Collectively, these data indicate that the narrow bud neck resulting from the same-site re-budding phenotype is unlikely a cause of a shorter RLS of *rga1*Δ mutants.

Since Rga1 plays a critical role in establishing the proper axis of Cdc42 polarization in G1 (Lee *et al*., 2015), we asked how Cdc42 polarization is affected in *rga1*Δ cells during aging. We imaged *rga1*Δ cells expressing PBD-RFP, which interacts specifically with Cdc42-GTP (Tong *et al*., 2007; Okada *et al*., 2017), and Cdc3-GFP (a septin subunit) as a marker for the cell division site and timing of the onset of cytokinesis. By quantifying the polarized PBD-RFP in young mother cells soon after the onset of cytokinesis, we found that the Cdc42-GTP level is about two-fold higher in *rga1*Δ cells than WT (Fig. 2A). Next, to monitor Cdc42 polarization in aged *rga1*Δ cells, we imaged old cells enriched by magnetic beads-based sorting (Sinclair, 2013). Interestingly, *rga1*Δ cells die predominantly at an unbudded state (94%, n = 52), unlike WT (46%, n = 50). These *rga1*Δ cells exhibited high PBD-RFP distribution over the entire mother cell cortex or in more than a single cluster near the terminal stage (Fig. 2, B & C; Fig. S1, a). These observations suggest that a defect in polarity establishment results in a shorter RLS of *rga1*Δ cells. Surprisingly, the Cdc42-GTP level was further elevated in the majority of *rga1*Δ cells before death (74.5%, n = 52 cells; Figs. 2B & S1, a), implying that hyperactivation of Cdc42 during aging involves additional mechanisms other than loss of Cdc42 GAP Rga1(see below).

**Figure 2.**
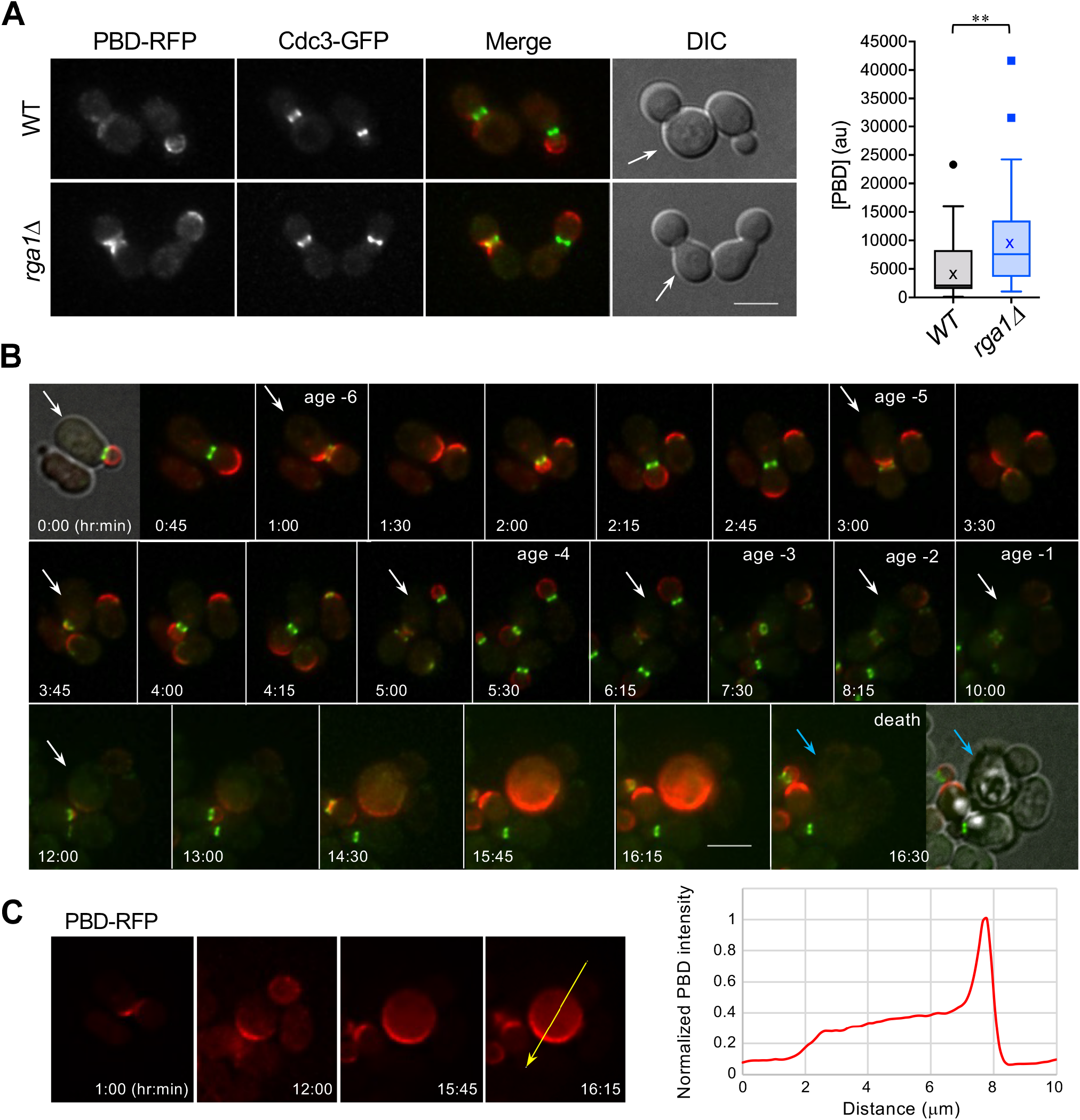
*rga1*Δ cells fail to establish polarity with a high Cdc42-GTP level near the terminal stage. **A**. Localization of PBD-RFP and Cdc3-GFP. Polarized Cdc42 during G1 was quantified in young mother cells of WT and *rga1*Δ (marked with arrows) by a threshold method (Okada *et al*., 2013): WT (n = 47) and *rga1*Δ (n = 53). In the box graph, quartiles and median values are shown with mean (x), and whiskers were created by the Tukey method. Scale bar, 5 µm. **B**. Time-lapse imaging of *rga1*Δ (expressing PBD-RFP and Cdc3-GFP) after mother cell sorting. The first and last images are shown as fluorescence images overlaid with DIC. Ages relative to the last cell division and selected timepoints (hr: min) are shown until death (marked with blue arrows). See also Fig. S1, a. **C**. Single Z slice images of PBD-RFP at selected timepoints (from Fig. 2B). Line scan shows PBD-RFP intensity along the arrow at 16:15, normalized to the peak intensity.

Suppression of a short RLS of *rga1*Δ mutants by *bud8*Δ (see Fig. 1E) suggests antagonistic roles of Bud8 and Rga1 in replicative aging. How does Bud8 impact RLS? Bud8 is known as a distal pole marker but is also observed at both poles of some newborn daughter cells (Taheri *et al*., 2000; Harkins *et al*., 2001). To gain insight into Bud8’s role in aging, we examined its localization in mother cells using mNeonGreen (mNG)-tagged Bud8. In addition to the bud tip, as previously reported, mNG-Bud8 localized to one pole or one side of the mother cell cortex similarly in **a** and **a**/α WT cells (Fig. 3A). Time-lapse imaging indicated that Bud8 at the bud tip is likely to remain at the distal pole in newborn cells after division, although the signal was often weaker in mother cells that had undergone multiple cell divisions (Fig. S2). Staining of the birth scar and bud scars confirmed that Bud8 localizes to the distal pole and weakly to the proximal pole (*i*.*e*., CRMs) in WT mother cells (Fig. 3A). Remarkably, Bud8 appeared in multiple puncta that overlapped with bud scars in WT mother cells that had divided several times (Figs. 3B & S2, b). This surprising localization pattern of Bud8 to multiple CRMs is reminiscent of that of Rax1 and Rax2 (Chen *et al*., 2000; Kang *et al*., 2004) and is consistent with Bud8’s interaction with Rax1 (Kang *et al*., 2004). Indeed, Bud8 localization was almost completely abolished at both poles of mother cells and diminished in buds in *rax1*Δ cells (Fig. 3, A & C). Time-lapse imaging showed that Bud8 exhibits significant colocalization with Rax2 in mother cells (Fig. S2, b & c). These results show that Bud8 is recruited to the distal pole and CRMs in mother cells by Rax1-Rax2.

**Figure 3.**
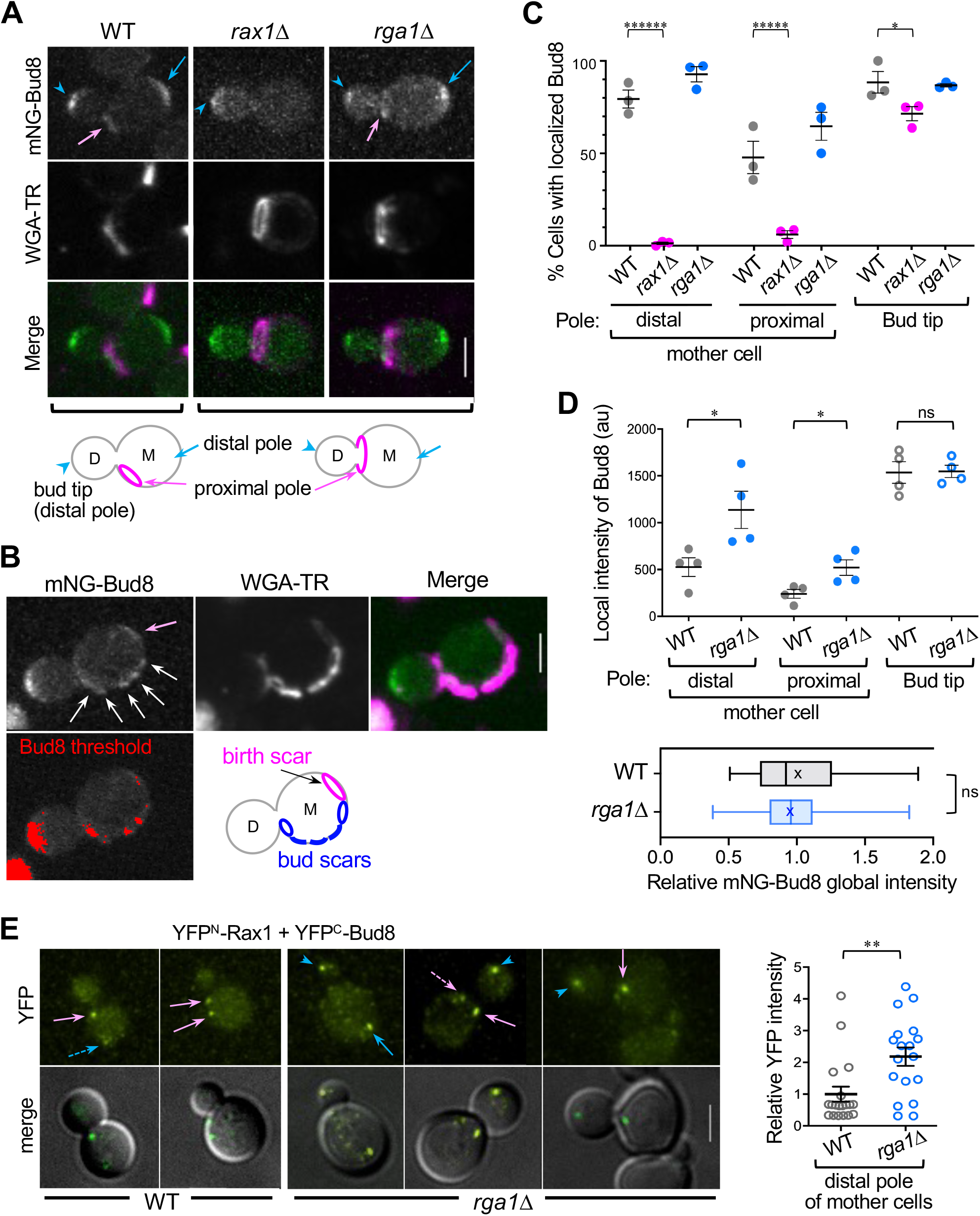
Bud8 localization and its interaction with Rax1. **A - B**. Localization of mNG-Bud8 in haploid cells stained with WGA-Texas red (TR). New mother cells with their first bud (**A**) and a mother cell that had budded several times (**B**) are shown. Bud8 signal at distinct locations are marked as depicted: the proximal poles (pink arrows), the distal pole of mother and bud (blue arrows and arrowheads), and bud scars (white arrows). The mNG-Bud8 intensity above threshold (**B)** is considered as localized signals. Scale bars, 3 µm. See also Fig. S2a. **C**. The percentage of cells with localized Bud8 at each site. Each symbol shows the mean value of an independent imaging set (n > 120 cells for each strain). Horizontal lines and error bars denote mean ± SEM, and *p* values are shown from unpaired two-tailed t-test. **D**. Quantification of the mNG-Bud8 localized to each site. Each symbol shows the mean value of an independent imaging set (n = 155, WT; n = 195 cells, *rga1*Δ) (see also legend to Fig. 3C). Below, relative global intensity of mNG-Bud8. Box graph shows quartiles and median values with mean (x), and whiskers show minimum to maximum values (n = 87, WT; n = 90 cells, *rga1*Δ). **E**. BiFC assay in WT and *rga1*Δ cells co-expressing YFP^N^-Rax1 and YFP^C^-Bud8. Pink and blue arrows mark BiFC signals at the proximal and distal pole, respectively, in mother cells; blue arrowheads mark the signal at the bud tip. Dotted arrows mark weak signals at each pole. Scale bar: 3 µm. Graph: The YFP intensity at the distal pole of individual mother cells is plotted: WT (n = 19) and *rga1*Δ (n = 18).

Since Rga1 is also recruited to CRMs (Miller *et al*., 2017), this localization pattern of Bud8 raises the possibility that Bud8 and Rga1 may compete for their localization to CRMs in mother cells. To test this idea, we examined Bud8 localization in *rga1*Δ cells. The number of cells with localized Bud8 signal in large-budded mother cells was slightly higher in *rga1*Δ cells than WT (Fig. 3, A & C). Interestingly, however, mNG-Bud8 levels at each pole of mother cells were significantly higher in *rga1*Δ than in WT cells, while the global intensity of mNG-Bud8 was similar in these cells (Fig. 3D). These results suggest that Bud8 is recruited to the distal pole and CRMs more efficiently in mother cells of *rga1*Δ than WT. To test this idea further, we examined *in vivo* interactions between Bud8 and Rax1 by bimolecular fluorescence complementation (BiFC) assays (Kerppola, 2008) using split YFP (yellow fluorescent protein) fragments fused to Bud8 and Rax1. As expected from their *in vitro* interaction and localization patterns (Kang *et al*., 2004)(see above), co-expression of YFP^N^-Rax1 and YFP^C^-Bud8 displayed YFP signals at CRMs and weakly at the distal pole of mother cells and bud tips (Fig. 3E). Because *rga1*Δ cells frequently re-bud at the previous bud site, each CRM is not separated as in WT, making it difficult to compare the YFP intensity at each CRM. We thus compared the YFP intensity at the distal pole of mother cells and found that the signal is higher in *rga1*Δ than WT cells (Fig. 3E). Although the BiFC signal at CRMs cannot be directly compared, this finding further supports the idea that Rga1 interferes with Bud8’s localization in mother cells.

Haploid WT cells bud in the axial pattern, and thus CRMs locate at or near the proximal pole in young mother cells. The interference of Bud8 localization to CRMs by Rga1 may account for their opposite impacts on RLS. However, Rga1 is not known to localize to the distal pole. How does Rga1 affect Bud8’s localization to the distal pole as well? This puzzle prompted us to re-examine Rga1 localization. Using a newly constructed, functional mNG-tagged Rga1, we found that mNG-Rga1 is enriched at the distal pole of mother cells and bud tips in addition to the bud neck and CRMs, albeit weaker and less tightly organized than its signal at the bud neck (Fig. S1, b). Comparison of the polarized mNG-Rga1 at the distal pole and CRMs in mother cells indicated that mNG-Rga1 was slightly more enriched at these sites in *bud8*Δ cells than WT, but the difference in young cells was not statistically significant (n = 85 – 112, *p* = 0.097; Fig. S1, b). Thus, Bud8 may not efficiently compete with Rga1 for its localization to the mother cells at least at young ages. Nonetheless, the impact of Rga1 on Bud8 localization and Bud8-Rax1 interaction (see Fig. 3) suggests that Rga1 hinders Bud8’s role by interfering with its localization to CRMs and the distal pole, although other explanations are possible. Understanding their counteracting roles in lifespan control will require further investigations, including how the localization of Rga1 or Bud8 changes during aging.

Because *rga1*Δ mother cells exhibit an elevation of Cdc42-GTP, particularly near the terminal stage (see Figs. 2B & S1, a), we wondered whether an excess Cdc42-GTP level could be a cause of senescence or aging, as reported in mammalian cells (Florian *et al*., 2012). We hypothesized that Cdc42-GTP accumulates in WT cells during aging (albeit at a lower extent than in *rga1*Δ cells) but much less in long-lived *bud8*Δ cells. To test this idea, we imaged the WT and *bud8*Δ cells expressing PBD-RFP and Cdc3-GFP in a microfluidic platform, which allows imaging of single cells until death by washing away daughter cells with continuous media flow (Xie *et al*., 2012; Chen *et al*., 2017).

By quantifying the PBD-RFP level during G1 (from the onset of cytokinesis just before bud emergence) at each age of individual WT cells, we observed stochastic fluctuation of Cdc42-GTP level during repeated cell divisions (70%, n = 40; Fig. 4A). The cross-section image of old WT cells indicated the enrichment of Cdc42-GTP at the cell cortex in addition to a higher concentration at a single polarized site (Fig. 4B). In contrast, *bud8*Δ cells rarely displayed such stochastic fluctuation of the Cdc42-GTP level in G1 during aging. Instead, old *bud8*Δ cells often showed PBD-RFP in the vacuolar lumen, which we confirmed by additional imaging with Vph1-GFP, a vacuole membrane marker (88%, n = 40; Fig. 4C). To avoid this signal in vacuoles, we quantified the PBD-RFP intensity along the mother cell periphery for comparison of the cortical Cdc42-GTP in WT and *bud8*Δ cells until death (see Materials and Methods). Each cell division was marked based on the peak PBD-RFP level in G1, and then single-cell trajectories were aligned with the last division of each cell (Fig. 4D). These analyses suggest that Cdc42-GTP rarely accumulates at the mother cell cortex in *bud8*Δ cells.

**Figure 4.**
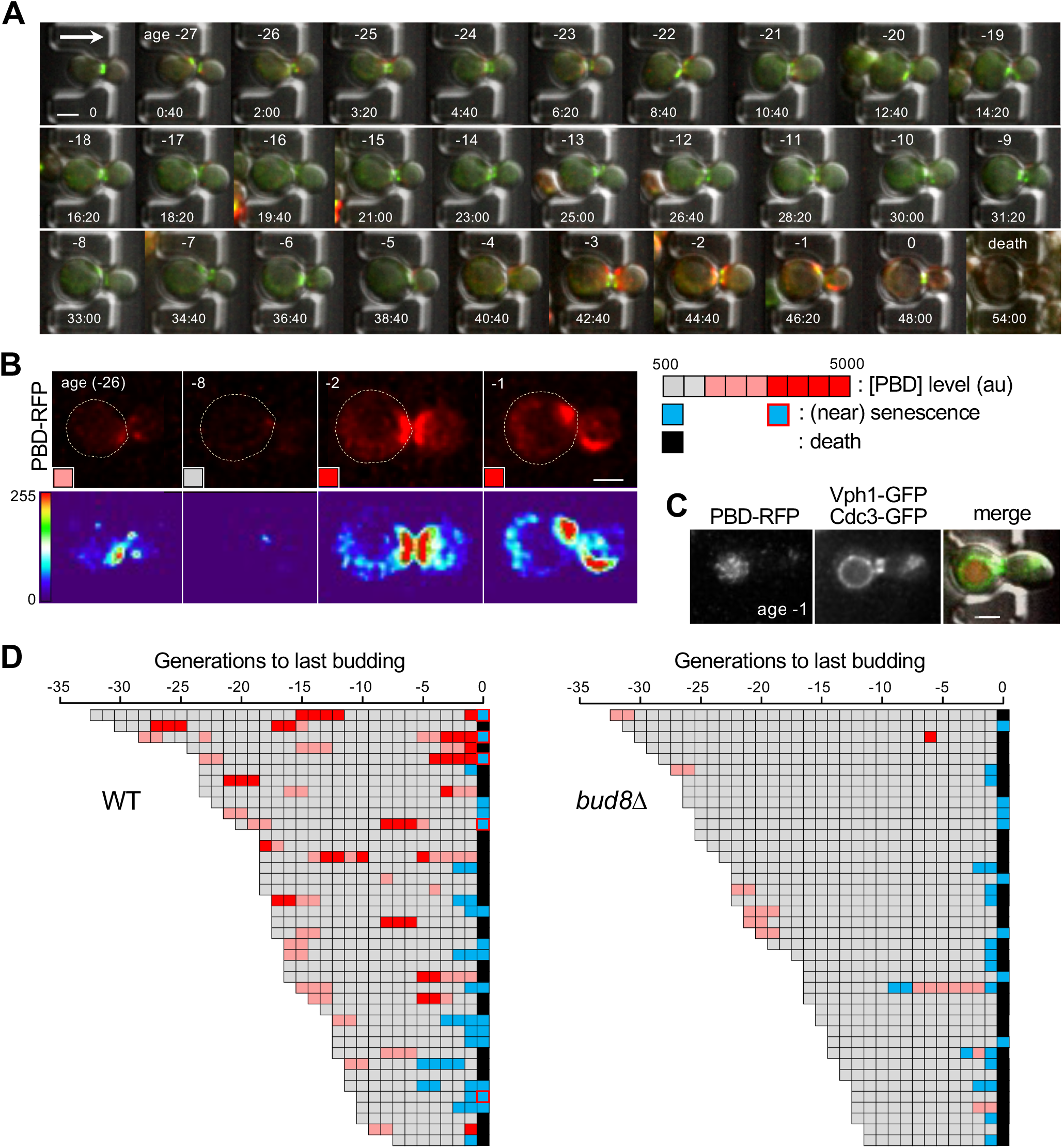
Cdc42-GTP is elevated in WT but rarely in *bud8*Δ cells. **A**. Microfluidic imaging of WT expressing PBD-RFP and Cdc3-GFP at 30 C. Cells around cytokinesis are shown, except the first initially loaded cell. Ages (relative to the last cell division) and timepoints (hr: min) are marked. Fluorescent images were deconvolved. Arrow marks the direction of media flow. Scale bar, 3 µm. **B**. Single Z slice images of PBD-RFP at selected timepoints (from Fig. 4A) are shown with heatmap histograms. Each image is marked with a colored box based on the peak PBD-RFP intensity along the cell periphery (see Materials and Methods). **C**. Localization of *bud8*Δ cells (*PBD-RFP CDC3-GFP*) carrying a Vph1-GFP plasmid. An old cell at the last division (age -1) is shown. **D**. Single-cell trajectories (each row) are aligned with the last cell division of WT and *bud8*Δ cells. Each cell division is marked based on the PBD-RFP level in G1 or death (see Fig. 4B). Divisions near or during senescence are scored as a sudden increase in the cell cycle length (> 5 hrs) with no clear sign of cell death; and are further distinguished with the PBD level as marked. Total numbers of cell divisions scored: n = 679 (WT) and n = 795 (*bud8*Δ) from 40 cells of each strain.

Our results described above support the idea that Rga1 and Bud8 may have opposite roles in aging by controlling the Cdc42-GTP level during repeated cell divisions. Although how aging impacts Rga1 or Bud8 is unknown, we considered a possibility that Rga1’s function may decline during aging. If Rga1’s localization to CRMs and the distal pole of mother cells decreases during aging, then artificially targeting Rga1 GAP persistently to the Rax1-Rax2 complex may extend lifespan. We tested this idea by fusing the Rga1 GAP domain (aa700-1007) to the C terminus of Rax2, which has type I membrane topology with its short C-terminal region in the cytoplasm (Kang *et al*., 2004). The expression of this Rax2-Rga1 GAP chimera improved the survival of mid-aged cells, slightly increasing RLS (median, 32; n = 92) compared to WT (median, 30; n = 48) (Fig. S1, c). However, its impact on older cells was negligible, and overall survival curves were not significantly different. We noted that a *bud8*Δ mutant also displays better survival than WT at mid ages but survives similarly to WT at old ages (see Fig. 1E). While the Rga1 GAP activity is necessary for longevity, RLS is thus likely to depend on the interplay between additional polarity factors such as Cdc42 (see below) and Bud8. This conclusion is also consistent with the observations of *rga1*Δ cells (Fig. 2). In addition, the *bud8*Δ *rga1*Δ double mutant has a similar RLS as WT rather than *rga1*Δ (Fig. 1E), suggesting that Bud8 may play an additional role in aging in addition to counteracting Rga1. Because of the low Cdc42-GTP accumulation at the mother cell cortex in *bud8*Δ cells, we speculate that Bud8 may recruit positive feedback of Cdc42 activation to the mother cell cortex during aging.

Our data discussed so far suggest that upregulation of Cdc42 activity during aging may lead to senescence and thus limit the lifespan. To test this idea further, we examined the impact of Cdc42 overexpression on RLS using a strain (*CDC42ov*) that carries extra copies of *GFP*-*CDC42* on the chromosome. This strain expresses GFP-Cdc42 moderately high – on average 2.2-fold higher than the *GFP-CDC42* strain with a single chromosomal copy of *GFP-CDC42* (Fig. 5A). This *CDC42ov* strain grew better than the *GFP-CDC42* strain (Wu *et al*., 2015) and even slightly faster than WT (with the endogenous, untagged *CDC42* gene) at 25 or 30°C (Fig. 5B). We performed microfluidic imaging of *GFP*-*CDC42ov* and WT (instead of the *GFP-CDC42* strain, which is a little sick at 30°C). The strains also expressed Cdc3-mCherry or Cdc3-GFP for efficient automated counting of cell divisions by a Python program (see Fig. S3; Materials and Methods). Strikingly, the *CDC42ov* strain underwent significantly fewer cell divisions than WT before death (Fig. 5C), indicating that overexpression of Cdc42 shortens RLS.

**Figure 5.**
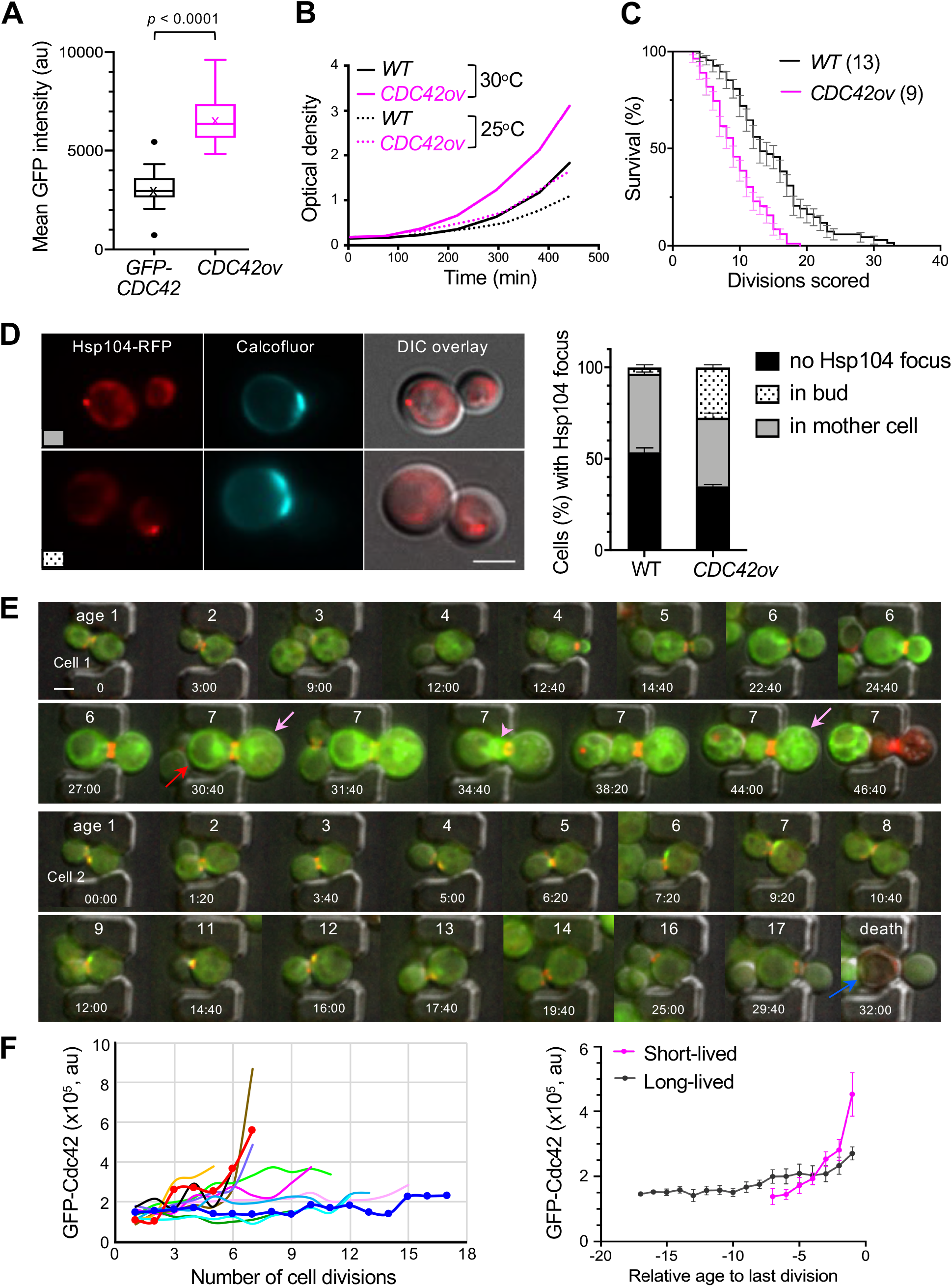
Mild overexpression of Cdc42 accelerates aging. **A**. GFP-Cdc42 levels in young cells (age 1) of *GFP-CDC42* (n = 24) and *GFP-CDC42ov* (n = 40). Box graph shows quartiles and median values with mean (x), and whiskers were created by the Tukey method. **B**. Growth curve of WT (with untagged *CDC42*) and *GFP-CDC42ov* strains. **C**. RLS estimation of WT (with untagged *CDC42* and *CDC3-GFP, n* = 67) and *GFP-CDC42ov* strain (n = 89) from microfluidic imaging at 30°C. The percentage (mean SEM) of surviving cells after each number of cell divisions is shown, and median RLS is indicated in parentheses. *p* < 0.0001, Log-rank (Mantel-Cox) test. These two strains are in the same background that is different from those used in Fig. 1. **D**. *CDC42ov* cells with Hsp104-RFP focus in mother (top) and daughter (bottom) cells. Hsp104-RFP often segregated to the bud in *CDC42ov* cells, confirmed by time-lapse imaging (not shown). The percentage (mean ± SEM) of cells with Hsp104 foci are shown on the graph by counting cells with medium-or large-size buds: WT (n = 110) and *CDC42ov* (n = 207). Scale bar, 3 μ m. **E**. Microfluidic images of *GFP-CDC42ov* (expressing Cdc3-mCherry) at 30°C. Cells at (near) cytokinesis are shown from age 1 with ages and time points (hr: min) marked. Fluorescent images were deconvolved. Cell 1 (representing short-lived cells) has likely entered senescence after the last division at age 7 (red arrow). The 7th daughter cell (pink arrow) initiated budding (pink arrowhead) died before completion of its first cell division. Cell 2 (representing long-lived cells) underwent asymmetric cell divisions until death at age 17. Scale bar, 3 μ m. **F**. Global intensity of GFP-Cdc42 during cytokinesis – G1 in *GFP-CDC42ov* cells at each age. Left: Colored lines represent 12 representative cells from age 1 to death or senescence. Red and blue lines with markers show cell 1 and 2 (see Fig. 5E), respectively. Right: GFP-Cdc42 level (mean SEM) is shown at each age (relative to the let division) for short-lived (less than 8 cell divisions) and longer-lived cells (n = 1 ∼ 24 cells at each age, each group).

Next, to test whether the shorter lifespan of the *CDC42ov* strain is indeed due to physiological aging, we examined the segregation of the Hsp104-associated focus in the *CDC42ov* cells. Aging is often associated with the formation of protein aggregates, which recruit the conserved heat-shock proteins (Hsps), such as Hsp104. Sequestration and asymmetric segregation of the protein aggregates are essential for the rejuvenation of asymmetrically dividing cells. Hsp104 functions as a chaperone in the recognition and disaggregation of the damaged proteins (Hill *et al*., 2017). As reported previously (Erjavec *et al*., 2007; Liu *et al*., 2010; Zhou *et al*., 2011; Saarikangas and Barral, 2015), an Hsp104-RFP focus (sometimes more than one Hsp104 foci) was faithfully retained in WT mother cells. In contrast, Hsp104 foci were often found in growing buds of the *Cdc42ov* at young ages (Fig. 5D). Such a defect has also been reported in *sir2*Δ cells (Erjavec *et al*., 2007), which lack Sir2, a conserved sirtuin anti-aging gene (Guarente, 2007). Collectively, these results suggest that mild overexpression of Cdc42 accelerates aging despite little impact on the fitness of young cells.

These *GFP-CDC4*2*ov* cells often exhibited distinct characteristics of naturally aging cells near senescence or death, e.g., enlarged, round cell shapes without Cdc42 enrichment at a single site, which suggest loss of cell polarity. Mothers sometimes produced daughter cells that were significantly larger in the last 20∼30% of their lifespan, resulting in daughters about the same size as (or even larger size than) the mother at the time of division (20%, n = 40; Fig. 5E). Symmetric cell division was also evident in old cells as newborn daughter cells died during their first few cell divisions. These characteristics of the *CDC4*2*ov* cells are remarkably similar to the previous observations on old yeast mother cells (Kennedy *et al*., 1994). We also noted that the GFP-Cdc42 level on the plasma membrane and endomembrane became quite heterogeneous among individual *CDC4*2*ov* cells as they aged, despite their similarity in young cells (at age 1) (Fig. 5F). Interestingly, cells with a shorter RLS had a higher accumulation of Cdc42 near the terminal stage than cells with a longer RLS (see Cell 1 and 2, Fig. 5E & Fig. 5F), implying the anti-correlation of the Cdc42 level and RLS. This accumulation of Cdc42 is likely to promote the elevation of Cdc42-GTP in aged cells, leading to the failure of polarity establishment. Consistent with this idea, mathematical modeling on spontaneous cell polarization predicted that a fraction of polarized cells decreases when the number of Cdc42 molecules in a cell increases (Altschuler *et al*., 2008).

Our data presented in this study suggest that upregulation of the activity and level of Cdc42 limits the RLS of budding yeast. While Cdc42 is essential for yeast cell proliferation, its elevation and hyperactivation may cause senescence in older cells by causing loss of cell polarity. The number of senescent cells increases with aging in multicellular organisms (Collado *et al*., 2007), while senescence is likely directly linked to RLS in yeast. Several studies in animals have suggested a causative role of Cdc42 in senescence or aging, although how it does so is unclear. Notably, aging is associated with increased Cdc42 activity, which correlates with the depolarized phenotype, symmetric cell divisions, and aging in hematopoietic stem cells and tissues in mice (Giebel, 2008; Florian *et al*., 2012; Florian *et al*., 2018). Inhibition of Cdc42 activity extends the lifespan of mice (Florian *et al*., 2020), while loss of a Cdc42 GAP shortens the lifespan of animals with premature aging phenotypes (Wang *et al*., 2007). Upregulation of Cdc42 is thus likely a general cause of senescence in asymmetrically dividing cells. The roles of Cdc42 and Bud8 in yeast cell polarity and aging are consistent with the ‘antagonistic pleiotropy’ hypothesis (Williams, 1957). This evolutionary theory of senescence posits that alleles may have opposite pleiotropic effects at different ages. If the beneficial effects of alleles early in life outweigh their deleterious effects at an old age, such genetic variants would be favored and enriched in a population (Williams, 1957; Gaillard and Lemaître, 2017). This study raises several unresolved questions, including how Cdc42 level increases in aging cells, how Bud8 promotes the elevation of cortical Cdc42-GTP during aging, and how aging impacts Rga1 or any other Cdc42 GAP. Yet it is tempting to speculate that a complex interplay among these polarity factors causes the stochastic elevation or activation of Cdc42, which may be one of the factors leading to the heterogeneity of lifespan among genetically identical cells in a population.

## MATERIALS AND METHODS

### Yeast strains and growth conditions

Standard methods of yeast genetics, DNA manipulation, and growth conditions were used (Guthrie and Fink, 1991). Yeast strains were grown in the appropriate synthetic medium containing 2% dextrose as a carbon source. Yeast strains used in this study are listed in Supplemental Table S1 with a brief description of construction methods. All yeast strains used for imaging express tagged proteins under their native promoters on the chromosomes, except Vph1-GFP, which was expressed from a CEN plasmid. For imaging, yeast strains were grown at 30°C, unless indicated otherwise, using the appropriate synthetic complete medium containing 2% dextrose as a carbon source. A ymNG (yeast mNeonGreen)-tagged Bud8 was expressed from the chromosomal locus of *BUD8* (see Table S1). ymNG-Bud8 was functional based on its budding pattern of an **a**/ cells (**a**/ *bud8* y*mNG-BUD8/ bud8* y*mNG-BUD8*) and localized similarly in **a** and **a**/ cells with an improved signal than GFP-Bud8.

### Determination of RLS

Lifespan analyses were carried out by using micromanipulation as described previously (Steffen *et al*., 2009) using YPD plates containing (2% [wt/vol] glucose and 2% [wt/vol] agar). During the intervening time of each micromanipulation, plates were sealed with parafilm and incubated at 30°C. Plates were shifted to 4°C for overnight incubation. To minimize any difference in growth conditions, lifespan assays of a mutant and the isogenic WT strain were performed in parallel. For all strains tested, mean RLS and *p* values were calculated from pooled experiments in which each strain of interest was compared with its respective WT strain. RLS was also estimated by counting the number of cell divisions from microfluidic images. A custom-made Python program was used for the analysis of properly oriented cells during imaging for the entire lifespan (see below). The number of cell divisions was manually counted with a visual inspection of microfluidic images of cells, whose positions were shifted relative to the pillars during imaging. Because of loss of full lifespan potential of daughter cells that are born from very old mothers (this study), consistent with a previous report (Kennedy *et al*., 1994), we excluded those cells that died in less than 3 cell divisions in determining RLS from microfluidic imaging data. The issue of daughters from very old mothers is negligible in a micromanipulation-based assay since such old mother cells are rare when cells are initially placed for the assay. However, such daughters can be trapped in the imaging platform in the middle of imaging because of their larger cell size, and thus the survival data can be skewed if these cells are included.

### Microscopy and image analysis

Cells were grown in an appropriate synthetic medium overnight and then freshly subcultured for 3–4 h in the same medium. For most time-lapse imaging, images were captured (9 z stacks, 0.4 μm step) every 10 or 15 min with cells either mounted on a 2% agarose slab or a glass-bottomed dish (MatTek) containing the indicated medium with 5 μM n-propyl gallate (Sigma), an anti-fade reagent, as previously described (Kang *et al*., 2018; Miller *et al*., 2019). The slab or dish was put directly on a stage (at 25–26°C) or in a temperature-control chamber set to 30°C, as indicated. Imaging involving mNG-Bud8 (Figs. 3 & S2) was performed using a spinning disk confocal microscope (Ultra-VIEW VoX CSU-X1 system; Perkin Elmer-Cetus) equipped with a 100x /1.45 NA Plan-Apochromat Lambda oil immersion objective lens (Nikon); 440-, 488-, 515- and 561-nm solid-state lasers (Modular Laser System 2.0; Perkin Elmer-Cetus); and a back-thinned EM CCD (ImagEM C9100-23B; Hamamatsu Photonics) on an inverted microscope (Ti-E; Nikon). Widefield fluorescence microscopy was performed for other live-cell imaging, including microfluidic imaging (see below), using an inverted microscope (Ti-E; Nikon) fitted with a 100x/1.45 NA Plan-Apochromat Lambda oil immersion objective lens (Nikon), FITC/GFP, and mCherry/Texas Red filters from Chroma Technology, an Andor iXon Ultra 888 electron-multiplying charge-coupled device (EM CCD) (Andor Technology), Sola Light Engine (Lumencor) solid-state illumination, and the software NIS Elements (Nikon).

To visualize the birth scar and bud scars in cells expressing mNG-Bud8, cells were stained with WGA-Texas Red (Thermo Fisher Scientific) at the final concentration of 100 µg/ml, and 15 z stacks (0.4 μm step) of static images were collected in both GFP and RFP channels and DIC (differential interference contrast), followed by time-lapse imaging only with GFP for mNG-Bud8 (Fig. S2, a) using a spinning disk confocal microscope (see above). Images of the birth scar and bud scars (Fig. 1F) were also captured similarly but using a Nikon Ti-E microscope (see above). Because of the distinct size difference, the birth scar (which is larger than bud scars) can be unambiguously identified from single-color imaging with WGA-Texas Red. Similarly, bud scars were visualized in cells expressing Hsp104-tdTomato (Fig. 5D), except staining cells with Calcofluor White (at 0.5 µg/ml). For time-lapse imaging of old mother cells until death, mother cells were enriched by a magnetic beads-based sorting protocol, as previously described (Sinclair, 2013), using EZ-Link Sulfo-NHS-LC-Biotin (Thermo Fisher Scientific) and ‘MACS cell separation’ reagents (Miltenyi Biotec), which include anti-biotin microbeads and LS columns (containing ferromagnetic sphere matrix). Mother cells enriched from the LS column were suspended in appropriate medium (containing propyl gallate) and then mounted in a low cell density in a glass-bottomed dish (see above), followed by time-lapse imaging for 8 ∼ 12 hrs at 30°C.

To make figures for fluorescence images, maximum intensity projections were generated using ImageJ (National Institutes of Health) or Fiji software for fluorescence images, and single z stack images were used for DIC images. Where indicated, images were deconvolved by the Iterative Constrained Richardson-Lucy algorithm using NIS Elements software for figure presentation. Image analyses were performed using ImageJ with summed intensity projections of z stacks after background subtraction. The Cdc42–GTP cluster in WT and *rga1* cells (Fig. 2A) was quantified by a threshold method using an ImageJ macro (Okada *et al*., 2017) from summed images of five selected z-sections after background subtraction, as previously described (Kang *et al*., 2014). For analysis of Bud8 localization pattern, a fluorescence threshold was set to select localized mNG-Bud8 clusters (see Fig. 3B), and then cells with or without the highlighted pixels at each location were counted (Fig. 3C). To measure the local intensity of mNG-Bud8, the same threshold was applied to all summed intensity projections, and integrated density was measured among all pixels above the threshold at each location.

Each data point in Fig. 3D represents an average of 4 independent imaging sets. To quantify global intensity of mNG-Bud8 (Fig. 3D), average intensity projections were created from all 13 z-sections (0.4-μm step), and an ROI was drawn around the outline of mother cells with large buds (by comparing the synchronized fluorescence and DIC channel images). WT cells without any fluorescently tagged protein were mixed to capture control cells together with experimental cells, and the mean intensity of the control cells was used as background. Global intensity of mNG-Bud8 in each cell was then calculated by subtracting background from the mean intensity of the whole cell. Global intensity of GFP-Cdc42 (Fig. 5, A & F) was quantified similarly, except in Fig. 5A, cells were pre-stained with Calcofluor White to select cells at age 1. For analysis of colocalization of mNG-Bud8 and Rax2-RFP in mother cells (Fig. S2, c), Pearson’s correlation coefficient was determined using ImageJ plugin JACoP (Bolte and Cordelières, 2006). To compare the localization of mNG-Bud8 and Rax2-tdTomato around the cell cortex in time-lapse images, freehand lines (3-pixel width, 0.432 µm-wide) were drawn along the cell periphery on single focused z-stack images using ImageJ, and the fluorescence intensities were then measured along the lines for each time point of time-lapse images (Fig. S2, b).

### Microfluidic imaging and image analysis

Microfluidics setup and growth conditions are essentially the same as described in (Singh *et al*., 2017). Microfluidic devices were fabricated using polydimethylsiloxane (PDMS) by adopting a design used in the Li laboratory (a kind gift from Rong Li). Yeast cells suspended at 10^6^–10^7^ cells/ml concentration were slowly loaded onto the device through the inlet until single cells were captured in features at the desired occupancy. Subsequently, fresh synthetic media was applied at the constant flow rate by a syringe pump (Fusion 200; Chemyx). An initial flow rate (30 μl/min) was applied to remove bubbles and wash out non-trapped cells, and then the flow was reduced to 10 μl/min to keep trapped mother cells under a positive pressure and to wash out daughter cells consistently. Time-lapse imaging was performed using an inverted widefield fluorescence microscope (Ti-E; Nikon) (see above) equipped with a 60x/1.4 NA Plan Apochromat Lambda oil immersion objective lens and DIC optics. Bright-field and fluorescence images were typically recorded for about 72 hrs at 20-min intervals using NIS Elements (Nikon). The temperature remained mostly constant at 29∼30°C during imaging, and time-lapse images at multiple XY positions were captured (5 z stacks, 0.5 μm step). Where indicated in figure legends, microfluidic images were deconvolved by the Iterative Constrained Richardson-Lucy algorithm (NIS Elements) and cropped at selected time points, preferentially at the cell division or soon after division to make figures. Images were adjusted with the same LUTs (look-up tables) setting.

Calcofluor-staining of bud scars indicated that the age of the WT and *bud8*Δ cells initially trapped in the microfluidics imaging chambers was about the same (average age = 1.2; n > 100).

Image analysis was performed after importing nd2 files using ImageJ Bio-Format importer plugin. Fluorescence intensity of PBD-RFP cluster in WT was compared by (i) quantifying the PBD-RFP cluster in summed images (of five z-sections) of a whole mother cells by a threshold method, as previously described (Okada *et al*. 2013; Okada *et al*., 2017), and (ii) by taking the mean value of the three peak pixel intensity from the PBD-RFP level along the mother cell periphery on single focused z-stack images using ImageJ freehand lines (3-pixel width, 0.644 µm-wide). Both methods indicated that the PBD-RFP level is the lowest at the onset of cytokinesis and then increases during G1 until bud emergence, as previously reported (Okada *et al*., 2013; Kang *et al*., 2014; Lee *et al*., 2015). Normalized values by both methods revealed similar fluctuation of Cdc42-GTP in G1 during repeated cell divisions, although the absolute values were different. However, because of the frequent appearance of PBD-RFP in the vacuolar lumen in old *bud8*Δ cells, the PBD-RFP data were likely to be skewed when summed z slice images were used for quantification of *bud8*Δ cells by the threshold method (i). Since Cdc42-GTP that was mis-targeted to vacuoles in *bud8*Δ cells is unlikely to contribute to Cdc42 polarization, the method (ii) was used to compare Cdc42-GTP cluster in WT and *bud8* cells (Fig. 4D). Even with the method (ii), there is a slight possibility of interference of the PBD-RFP signal from vacuoles because of the proximity of the large vacuole (in the aged cells) to the plasma membrane with the resolution of typical microfluidic images. For this reason, the PBD-RFP levels during G1 in every cell division are marked in three groups in Fig. 4D, as marked in Fig. 4B (instead of displaying the exact PBD level in each G1). Cell death was evident morphologically with abruptly shrinking cell size or cell lysis (from DIC images), which was often accompanied with sudden complete loss of normal fluorescence signals or abrupt appearance of strong autofluorescence in the whole cell in both channels. Cell divisions are marked as senescence or near senescence when there was a sudden increase in the cell cycle length (> 5 ∼ 6 hrs) with no clear evidence of cell death and when no further division was scored within 8 ∼ 12 hrs. To compare the GFP-Cdc42 level in a strain carrying a single copy (*GFP-CDC42*) or *GFP-Cdc42ov*, the mean global intensity of GFP was quantified as described above. To quantify the GFP-Cdc42 level during aging, the mean intensity of GFP-Cdc42 was measured similarly soon after the onset of cytokinesis or G1 at each age until death or senescence. Total GFP-Cdc42 level at each age was calculated by multiplying mean intensity by cell size at each age in individual cells (n = 40). Total GFP-Cdc42 level at each age was then plotted for 12 representative cells with absolute ages (Fig. 5F, graph on the left). The data are also grouped based on the RLS of each cell – short-lived (less than 8 cell divisions in total) and long-lived group. The total GFP-Cdc42 level (mean ± SEM) at each age was then plotted with respect to the relative age to the last division (Fig. 5F, graph on the right).

### Program for RLS measurement

We developed a Python program for automatic analysis of RLS of multiple single cells in one XY position of capturing. This program allows rapid RLS determination based on fluctuation of a fluorescent marker level during the cell cycle (such as Cdc3-GFP or Cdc3-mCherry marker, whose level drops in the G1 phase when the old ring disassembles). Because this analysis identifies each cell based on its position, the program works well for those cells whose positions within the PDMS pillars have not shifted much during the entire lifespan. RLS measurement of a cell expressing Cdc3-GFP (and PBD-RFP) by the program and ImageJ is compared (see an example in Fig. S3, f). The automatic reading of the Cdc3-GFP level of each capture showed a repeating pattern of a curve, and the number of the valleys on the graph matches the number of cell divisions until death. However, the current program has some limitation because some cells rotate during imaging or shift their positions further away from the PDMS pillars (while these cells remain trapped). The algorithm and the detailed steps of this program are described below.

Cropping the whole chip:

Since the whole microfluidics chip has a specific layout of chambers (*i*.*e*., 4 columns per XY position; and 5 or 6 chambers per column; see Fig. S2, a), this provides a convenient way to crop an XY capture into multiple images covering only one cell in a pair of pillars. Cropping of images can be done by first cutting the whole capture vertically into columns then cutting horizontally. This is done by calculating the average intensity in a small sliding window through the whole image first horizontally and then vertically to determine the cut points (Fig. S2).

Detailed algorithm steps:

1. Load the nd2 image file with BioFormats function from pims library.
2. Keep the DIC channel from the loaded file for cropping.
3. Define a square sliding window with size 1 ^*^ max_height
4. Collect all the average intensity from x = 0 to x = max_width
5. Find the max intensity position *p_1* from average intensities. Add *p_1* to horizontal cutting points.
6. Set all the average intensity values to 0 and cut one column of chambers out (see Fig. S2, b & c) if its position *abs(p_2-p_1) < T*. T is a threshold which should be set slightly larger than the chamber width.
7. Repeat step 5 and 6 for 4 times.
8. Define a square sliding window with size column_width ^*^1
9. Collect all the average intensity from y = 0 to y = max_width
10. Similar to step 5 and 6, pick the vertical cutting points and crop each chamber out.
11. Repeat step 10 for 6 times if the column is the first and the third to the left, otherwise 5 times.
12. Return the cropping results.

### Statistical analysis

Data analysis was performed using Prism 8 (GraphPad Software). To determine statistical differences between two sets of data of cell survival, the Gehan-Breslow-Wilcoxon test and Log-rank (Mantel-Cox) test were used. Where indicated in Figure legends, a two-tailed student’s t test was performed to determine statistical significance: ns (not significant) for p ≥ 0.05, ^*^p < 0.05, ^**^p < 0.01, ^***^p < 0.001, ^****^p < 0.0001, ^*****^*p* < 0.00001, and ^******^*p* < 0.000001. Horizontal lines and error bars on the graphs (scattered plots) denote mean ± SEM. In the box graphs, quartiles and median values are shown together with mean (marked with x). Unless indicated otherwise, the Tukey method was used to create the whiskers, which indicate variability outside the upper and lower quartiles. Any point outside those whiskers is considered an outlier.

## Abbreviations

(CRMs): cytokinesis remnants
(RLS): replicative lifespan
(GAP): GTPase-activating protein
(WT): wild type
WGA: (Wheat germ agglutinin)

## Acknowledgments

We are grateful to Rong Li (Johns Hopkins) and Hao Li (UCSF) for their help with microfluidics and K. E. Miller for her assistance with microfluidic imaging. We also thank E. Bi, J. R. Pringle, and D. Lew for yeast strains and plasmids. This work has been supported by grants from the National Institutes of Health/National Institute of General Medical Sciences (R01-GM114582 to H.-O. P) and the NIH/National Institute on Aging (R21-AG060028 to H.-O. P. & P. J. K). The authors declare no competing financial interests.

## Supplemental Material

Figure S1 – S3

Table S1

## Notes

### Competing Interest Statement

The authors have declared no competing interest.

### Summary of Updates

New figures are added (Fig. 2 & Fig. S1). Fig 1 - 5 are revised.

## REFERENCES

Altschuler, S.J., Angenent, S.B., Wang, Y., and Wu, L.F. (2008). On the spontaneous emergence of cell polarity. Nature 454, 886–889.

Barton, A.A. (1950). Some aspects of cell division in saccharomyces cerevisiae. J. Gen. Microbiol. 4, 84–86.

Bi, E., and Park, H.-O. (2012). Cell polarization and cytokinesis in budding yeast. Genetics 191, 347–387.

Bolte, S., and Cordelières, F.P. (2006). A guided tour into subcellular colocalization analysis in light microscopy. J. Microsc. 224, 213–232.

Chen, K.L., Crane, M.M., and Kaeberlein, M. (2017). Microfluidic technologies for yeast replicative lifespan studies. Mech. Ageing Dev. 161, 262–269.

Chen, T., Hiroko, T., Chaudhuri, A., Inose, F., Lord, M., Tanaka, S., Chant, J., and Fujita, A. (2000). Multigenerational cortical inheritance of the Rax2 protein in orienting polarity and division in yeast. Science 290, 1975–1978.

Collado, M., Blasco, M.A., and Serrano, M. (2007). Cellular senescence in cancer and aging. Cell 130, 223–233.

Egilmez, N.K., and Jazwinski, S.M. (1989). Evidence for the involvement of a cytoplasmic factor in the aging of the yeast Saccharomyces cerevisiae. J. Bacteriol. 171, 37–42.

Erjavec, N., Larsson, L., Grantham, J., and Nyström, T. (2007). Accelerated aging and failure to segregate damaged proteins in Sir2 mutants can be suppressed by overproducing the protein aggregation-remodeling factor Hsp104p. Genes Dev. 21, 2410–2421.

Florian, M.C., Dorr, K., Niebel, A., Daria, D., Schrezenmeier, H., Rojewski, M., Filippi, M.D., Hasenberg, A., Gunzer, M., Scharffetter-Kochanek, K., Zheng, Y., and Geiger, H. (2012). Cdc42 activity regulates hematopoietic stem cell aging and rejuvenation. Cell Stem Cell 10, 520–530.

Florian, M.C., Klose, M., Sacma, M., Jablanovic, J., Knudson, L., Nattamai, K.J., Marka, G., Vollmer, A., Soller, K., Sakk, V., Cabezas-Wallscheid, N., Zheng, Y., Mulaw, M.A., Glauche, I., and Geiger, H. (2018). Aging alters the epigenetic asymmetry of HSC division. PLoS biology 16, e2003389.

Florian, M.C., Leins, H., Gobs, M., Han, Y., Marka, G., Soller, K., Vollmer, A., Sakk, V., Nattamai, K.J., Rayes, A., Zhao, X., Setchell, K., Mulaw, M., Wagner, W., Zheng, Y., and Geiger, H. (2020). Inhibition of Cdc42 activity extends lifespan and decreases circulating inflammatory cytokines in aged female C57BL/6 mice. Aging Cell 19, e13208.

Gaillard, J.M., and Lemaître, J.F. (2017). The Williams’ legacy: A critical reappraisal of his nine predictions about the evolution of senescence. Evolution 71, 2768–2785.

Giebel, B. (2008). Cell polarity and asymmetric cell division within human hematopoietic stem and progenitor cells. Cells Tissues Organs 188, 116–126.

Guarente, L. (2007). Sirtuins in aging and disease. Cold Spring Harb. Symp. Quant. Biol. 72, 483–488.

Guthrie, C., and Fink, G.R. (1991). Guide to Yeast Genetics and Molecular Biology. Academic Press: San Diego.

Harkins, H.A., Pagé, N., Schenkman, L.R., De Virgilio, C., Shaw, S., Bussey, H., and Pringle, J.R. (2001). Bud8p and Bud9p, proteins that may mark the sites for bipolar budding in yeast. Mol. Biol. Cell 12, 2497–2518.

Hill, S.M., Hanzén, S., and Nyström, T. (2017). Restricted access: spatial sequestration of damaged proteins during stress and aging. EMBO Rep 18, 377–391.

Jazwinski, S.M., Kim, S., Lai, C.Y., and Benguria, A. (1998). Epigenetic stratification: the role of individual change in the biological aging process. Exp. Gerontol. 33, 571–580.

Johnston, J.R. (1966). Reproductive capacity and mode of death of yeast cells. Antonie Van Leeuwenhoek 32, 94–98.

Kang, P.J., Angerman, E., Nakashima, K., Pringle, J.R., and Park, H.-O. (2004). Interactions among Rax1p, Rax2p, Bud8p, and Bud9p in marking cortical sites for bipolar bud-site selection in yeast. Mol. Biol. Cell 15, 5145–5157.

Kang, P.J., Lee, M.E., and Park, H.-O. (2014). Bud3 activates Cdc42 to establish a proper growth site in budding yeast. J. Cell Biol. 206, 19–28.

Kang, P.J., Miller, K.E., Guegueniat, J., Beven, L., and Park, H.-O. (2018). The shared role of the Rsr1 GTPase and Gic1/Gic2 in Cdc42 polarization. Mol. Biol. Cell 29, 2359–2369.

Kennedy, B.K., Austriaco, N.R., Jr., and Guarente, L. (1994). Daughter cells of Saccharomyces cerevisiae from old mothers display a reduced life span. J. Cell Biol. 127, 1985–1993.

Kenyon, C. (2001). A conserved regulatory system for aging. Cell 105, 165–168.

Kerppola, T.K. (2008). Bimolecular fluorescence complementation (BiFC) analysis as a probe of protein interactions in living cells. Annu Rev Biophys 37, 465–487.

Lee, M.E., Lo, W.C., Miller, K.E., Chou, C.S., and Park, H.-O. (2015). Regulation of Cdc42 polarization by the Rsr1 GTPase and Rga1, a Cdc42 GTPase-activating protein, in budding yeast. J. Cell Sci. 128, 2106–2117.

Liu, B., Larsson, L., Caballero, A., Hao, X., Oling, D., Grantham, J., and Nyström, T. (2010). The polarisome is required for segregation and retrograde transport of protein aggregates. Cell 140, 257–267.

McCormick, M.A., Delaney, J.R., Tsuchiya, M., Tsuchiyama, S., Shemorry, A., Sim, S., Chou, A.C., Ahmed, U., Carr, D., Murakami, C.J., Schleit, J., Sutphin, G.L., Wasko, B.M., Bennett, C.F., Wang, A.M., Olsen, B., Beyer, R.P., Bammler, T.K., Prunkard, D., Johnson, S.C., Pennypacker, J.K., An, E., Anies, A., Castanza, A.S., Choi, E., Dang, N., Enerio, S., Fletcher, M., Fox, L., Goswami, S., Higgins, S.A., Holmberg, M.A., Hu, D., Hui, J., Jelic, M., Jeong, K.S., Johnston, E., Kerr, E.O., Kim, J., Kim, D., Kirkland, K., Klum, S., Kotireddy, S., Liao, E., Lim, M., Lin, M.S., Lo, W.C., Lockshon, D., Miller, H.A., Moller, R.M., Muller, B., Oakes, J., Pak, D.N., Peng, Z.J., Pham, K.M., Pollard, T.G., Pradeep, P., Pruett, D., Rai, D., Robison, B., Rodriguez, A.A., Ros, B., Sage, M., Singh, M.K., Smith, E.D., Snead, K., Solanky, A., Spector, B.L., Steffen, K.K., Tchao, B.N., Ting, M.K., Vander Wende, H., Wang, D., Welton, K.L., Westman, E.A., Brem, R.B., Liu, X.G., Suh, Y., Zhou, Z., Kaeberlein, M., and Kennedy, B.K. (2015). A Comprehensive Analysis of Replicative Lifespan in 4,698 Single-Gene Deletion Strains Uncovers Conserved Mechanisms of Aging. Cell Metab. 22, 895–906.

Meitinger, F., Khmelinskii, A., Morlot, S., Kurtulmus, B., Palani, S., Andres-Pons, A., Hub, B., Knop, M., Charvin, G., and Pereira, G. (2014). A memory system of negative polarity cues prevents replicative aging. Cell 159, 1056–1069.

Miller, K.E., Lo, W.C., Chou, C.S., and Park, H.-O. (2019). Temporal regulation of cell polarity via the interaction of the Ras GTPase Rsr1 and the scaffold protein Bem1. Mol. Biol. Cell 30, 2543–2557.

Miller, K.E., Lo, W.C., Lee, M.E., Kang, P.J., and Park, H.-O. (2017). Fine-tuning the orientation of the polarity axis by Rga1, a Cdc42 GTPase-activating protein. Mol. Biol. Cell 28, 3773–3788.

Mortimer, R.K., and Johnston, J.R. (1959). Life span of individual yeast cells. Nature 183, 1751–1752.

Nestelbacher, R., Laun, P., and Breitenbach, M. (1999). Images in experimental gerontology. A senescent yeast mother cell. Exp. Gerontol. 34, 895–896.

Okada, S., Leda, M., Hanna, J., Savage, N.S., Bi, E., and Goryachev, A.B. (2013). Daughter cell identity emerges from the interplay of Cdc42, septins, and exocytosis. Dev. Cell 26, 148–161.

Okada, S., Lee, M.E., Bi, E., and Park, H.-O. (2017). Probing Cdc42 Polarization Dynamics in Budding Yeast Using a Biosensor. Methods Enzymol. 589, 171–190.

Saarikangas, J., and Barral, Y. (2015). Protein aggregates are associated with replicative aging without compromising protein quality control. eLife 4, e06197.

Sinclair, D.A. (2013). Studying the Replicative Life Span of Yeast Cells. Methods in molecular biology (Clifton, N.J.) 1048, 49–63.

Singh, P., Ramachandran, S.K., Zhu, J., Kim, B.C., Biswas, D., Ha, T., Iglesias, P.A., and Li, R. (2017). Sphingolipids facilitate age asymmetry of membrane proteins in dividing yeast cells. Mol. Biol. Cell 28, 2712–2722.

Smith, G.R., Givan, S.A., Cullen, P., and Sprague, G.F., Jr. (2002). GTPase-activating proteins for Cdc42. Eukaryotic Cell 1, 469–480.

Steffen, K.K., Kennedy, B.K., and Kaeberlein, M. (2009). Measuring replicative life span in the budding yeast. J Vis Exp.

Steinkraus, K.A., Kaeberlein, M., and Kennedy, B.K. (2008). Replicative aging in yeast: the means to the end. Annual review of cell and developmental biology 24, 29–54.

Taheri, N., Köhler, T., Braus, G.H., and Mösch, H.-U. (2000). Asymmetrically localized Bud8p and Bud9p proteins control yeast cell polarity and development. The EMBO J. 19, 6686–6696.

Tong, Z., Gao, X.D., Howell, A.S., Bose, I., Lew, D.J., and Bi, E. (2007). Adjacent positioning of cellular structures enabled by a Cdc42 GTPase-activating protein-mediated zone of inhibition. J. Cell Biol. 179, 1375–1384.

Wang, L., Yang, L., Debidda, M., Witte, D., and Zheng, Y. (2007). Cdc42 GTPase-activating protein deficiency promotes genomic instability and premature aging-like phenotypes. Proceedings of the National Academy of Sciences of the United States of America 104, 1248–1253.

Wasko, B.M., and Kaeberlein, M. (2014). Yeast replicative aging: a paradigm for defining conserved longevity interventions. FEMS Yeast Res 14, 148–159.

Williams, G.C. (1957). Pleiotropy, natural selection, and the evolution of senescence. Evolution 11, 398–411.

Wu, C.F., Chiou, J.G., Minakova, M., Woods, B., Tsygankov, D., Zyla, T.R., Savage, N.S., Elston, T.C., and Lew, D.J. (2015). Role of competition between polarity sites in establishing a unique front. eLife 4, 11611.

Xie, Z., Zhang, Y., Zou, K., Brandman, O., Luo, C., Ouyang, Q., and Li, H. (2012). Molecular phenotyping of aging in single yeast cells using a novel microfluidic device. Aging Cell 11, 599–606.

Zahner, J.E., Harkins, H.A., and Pringle, J.R. (1996). Genetic analysis of the bipolar pattern of bud-site selection in the yeast Saccharomyces cerevisiae. Mol. Cell. Biol. 16, 1857–1870.

Zhou, C., Slaughter, B.D., Unruh, J.R., Eldakak, A., Rubinstein, B., and Li, R. (2011). Motility and segregation of Hsp104-associated protein aggregates in budding yeast. Cell 147, 1186–1196.

