## Supplemental Table & Figures for "Upregulation of the Cdc42 GTPase limits the replicative lifespan of budding yeast"

**Table S1. Yeast strains used in this study**

| Strain | Relevant Genotype[a] | Source/Comments |
|---|---|---|
| BY4741[@] | *MAT***a** *his3-Δ1 leu2Δ0 met15Δ0 ura3Δ0* | Open Biosystems |
| HPY2206[@] | *MAT***a** *rga1Δ::KAN* | Open Biosystems |
| HPY2438[@] | *MAT***a** *bud8Δ::KAN* | Open Biosystems |
| HPY3700[@] | *MAT***a** *bud8Δ::KAN rga1Δ::KAN* | This study |
| HPY2247[@] | *MAT***a** *rga2Δ::KAN* | Open Biosystems |
| HPY2425[@] | *MAT***a** *bem3Δ::KAN* | Open Biosystems |
| HPY664[@] | *MAT***a** *rsr1Δ::KAN* | Open Biosystems |
| HPY1187[@] | *MAT***a** *axl1Δ::KAN* | Open Biosystems |
| HPY1444[@] | *MAT***a** *axl2Δ::KAN* | Open Biosystems |
| HPY2681[#] | *MAT***a** *CDC3-GFP-LEU2* | This study[b] |
| HPY2325[#] | *MAT***a** *GIC2-PBD-tdTomato-URA3 CDC3-GFP-LEU2* | This study[c] |
| HPY3469[#] | *MAT***a** *bud8Δ::TRP1 GIC2-PBD-tdTomato-URA3 CDC3-GFP-LEU2* | This study |
| HPY3487[#] | *MAT***a** *bud8Δ::KAN GIC2-PBD-tdTomato-URA3 CDC3-GFP-LEU2* | This study |
| HPY3586[#] | *MAT***a** *bud8Δ::TRP1* y*mNG-BUD8-URA3* | This study[d] |
| HPY3592[#] | *MAT***a**/*MAT*α *bud8Δ::TRP1/ bud8Δ::TRP1* y*mNG-BUD8-URA3/* y*mNG-BUD8-URA3* | This study |
| HPY3671[#] | *MAT***a** *bud8Δ::TRP1* y*mNG-BUD8-URA3 rax1Δ::HIS3* | This study |
| HPY3672[#] | *MAT***a** *bud8Δ::TRP1* y*mNG-BUD8-URA3 rga1Δ::HIS3* | This study |
| HPY3694[#] | *MAT***a** *rga1Δ::HIS3 GIC2-PBD-tdTomato-URA3 CDC3-GFP-LEU2* | This study |
| HPY3702[#] | *MAT***a** *bud8Δ::TRP1* y*mNG-BUD8-URA3 RAX2-tdTomato-KAN* | This study |
| HPY3721[#] | *MAT***a** *cdc42::TRP1 GFP-CDC42(8X)-URA3 CDC3-mCherry-LEU2* | This study[e] |
| HPY3733[#] | *MAT***a** *cdc42Δ::TRP1 GFP-CDC42(1X)-URA3 CDC3-mCherry-LEU2* | This study[f] |
| HPY3748[#] | *MAT***a** *cdc42::TRP1 GFP-CDC42(8X)-URA3 HSP104-tdTomato-KAN* | This study |
| HPY3750[#] | *MAT***a** *HSP104-tdTomato-KAN* | This study[g] |
| HPY3764[#] | *MAT*α *rga1Δ::HIS3* y*mNG-RGA1-URA3* | This study[h] |
| HPY3794[#] | *MAT***a** *rga1Δ::HIS3* y*mNG-RGA1-URA3 bud8Δ::TRP1* | This study |

| | | |
|---|---|---|
| HPY3771[#] | *MAT***a** *bud8Δ::TRP1 YFP*$^{C}$*-BUD8-URA3 rax1Δ::URA3 YFP*$^{N}$*-RAX1-HIS3* | This study[i] |
| HPY3780[#] | *MAT*α *bud8Δ::TRP1 YFP*$^{C}$*-BUD8-URA3 rax1Δ::URA3 YFP*$^{N}$*-RAX1-HIS3 rga1Δ::HIS3* | This study |
| HPY3797[@] | *MAT***a** *rax2Δ::HIS3 RAX2-RGA1-GAP-LEU2* | This study[j] |

[a] Strains marked with [#] are derived from YEF473 (Bi and Pringle, 1996); strains marked with [@] are derived BY4741 (in S288C background).

[b] YIplac128-CDC3-GFP (a gift from E. Bi) (Tong *et al.*, 2007) was linearized with *Bgl*II and integrated into the *CDC3* locus.

[c] YIp211-GIC2-PBD-tdTomato carrying Gic2(1-208)-*tdTomato* (a gift from E. Bi) (Tong *et al.*, 2007) was linearized with *Apa*I and integrated into the *URA3* locus.

[d] YIp211-ymNG-BUD8 (pHP2274) was linearized by digesting with *Bam*HI and then integrated into the *BUD8* locus. To fuse ymNG (yeast mNeonGreen) to the N-terminus of Bud8, YIp211-ymNG-BUD8 (pHP2274) was generated by replacing the HA epitope sequence of YIp211-HA-BUD8 (a gift from J. Pringle) with the *Not*I cassette carrying the ymNG coding sequence derived from pFA6a-ymNeonGreen-CaURA3 (Addgene #125703).

[e] Derived from DLY13891 (a gift from D. Lew) (Wu *et al.*, 2015).

[f] YIplac211-$P_{CDC42}$-GFP-linker-CDC42 (a gift from D. Lew) was linearized with *EcoR*V and integrated into the *URA3* locus.

[g] A standard PCR-mediated tagging of tdTomato to the C terminus of Hsp104 was performed using pFA6a-tdTomato-KAN as a template.

[h] YIp211-ymNG-RGA1 (pHP2280) was linearized by digesting with *Apa*I and then integrated into the *URA3* locus. YIp211-ymNG-RGA1 was generated by replacing the HA epitope sequence of YIp211-HA-RGA1 (a gift from E. Bi) (Caviston *et al.*, 2003) with the *Not*I cassette carrying the ymNG coding sequence.

[i] YIp211-YFP$^{C}$-BUD8 (pHP2278) was linearized by digesting with *Bam*HI and then integrated into the *BUD8* locus. To fuse YFP$^{C}$ to the N-terminus of Bud8, the ymNG sequence of YIp211-ymNG-BUD8 (pHP2274) was replaced with the *Not*I cassette carrying the YFP$^{C}$ (Singh *et al.*, 2008). To fuse YFP$^{N}$ to the N-terminus of Rax1, pRS303-YFP$^{N}$-RAX1 (pHP1626), which was derived from pHP1061 (Kang *et al.*, 2004), was linearized by digesting with B*sp*MI and then integrated into the *RAX1* locus.

[j] *Bam*HI cassette of PCR fragment encoding GAP domain of Rga1 (aa 700-1007) was inserted in frame at a *Bam*HI site of *RAX2* (at aa 977, just after the transmembrane domain), and the resulting plasmid (pHP2289) was linearized by *Hind*III digestion and integrated at the *RAX2* chromosomal site.

**References for Supplemental Table S1**


Bi, E., and Pringle, J.R. (1996). *ZDS1* and *ZDS2*, genes whose products may regulate Cdc42p in *Saccharomyces cerevisiae*. Mol. Cell. Biol. *16*, 5264-5275.

Caviston, J.P., Longtine, M., Pringle, J.R., and Bi, E. (2003). The role of Cdc42p GTPase-activating proteins in assembly of the septin ring in yeast. Mol. Biol. Cell *14*, 4051-4066.

Kang, P.J., Angerman, E., Nakashima, K., Pringle, J.R., and Park, H.-O. (2004). Interactions among Rax1p, Rax2p, Bud8p, and Bud9p in marking cortical sites for bipolar bud-site selection in yeast. Mol. Biol. Cell *15*, 5145-5157.

Singh, K., Kang, P.J., and Park, H.-O. (2008). The Rho5 GTPase is necessary for oxidant-induced cell death in budding yeast. Proc. Natl. Acad. Sci. USA. *105*, 1522-1527.

Tong, Z., Gao, X.D., Howell, A.S., Bose, I., Lew, D.J., and Bi, E. (2007). Adjacent positioning of cellular structures enabled by a Cdc42 GTPase-activating protein-mediated zone of inhibition. J. Cell Biol. *179*, 1375-1384.

Wu, C.F., Chiou, J.G., Minakova, M., Woods, B., Tsygankov, D., Zyla, T.R., Savage, N.S., Elston, T.C., and Lew, D.J. (2015). Role of competition between polarity sites in establishing a unique front. eLife *4*, 11611.

**SUPPLEMENTAL FIGURES**

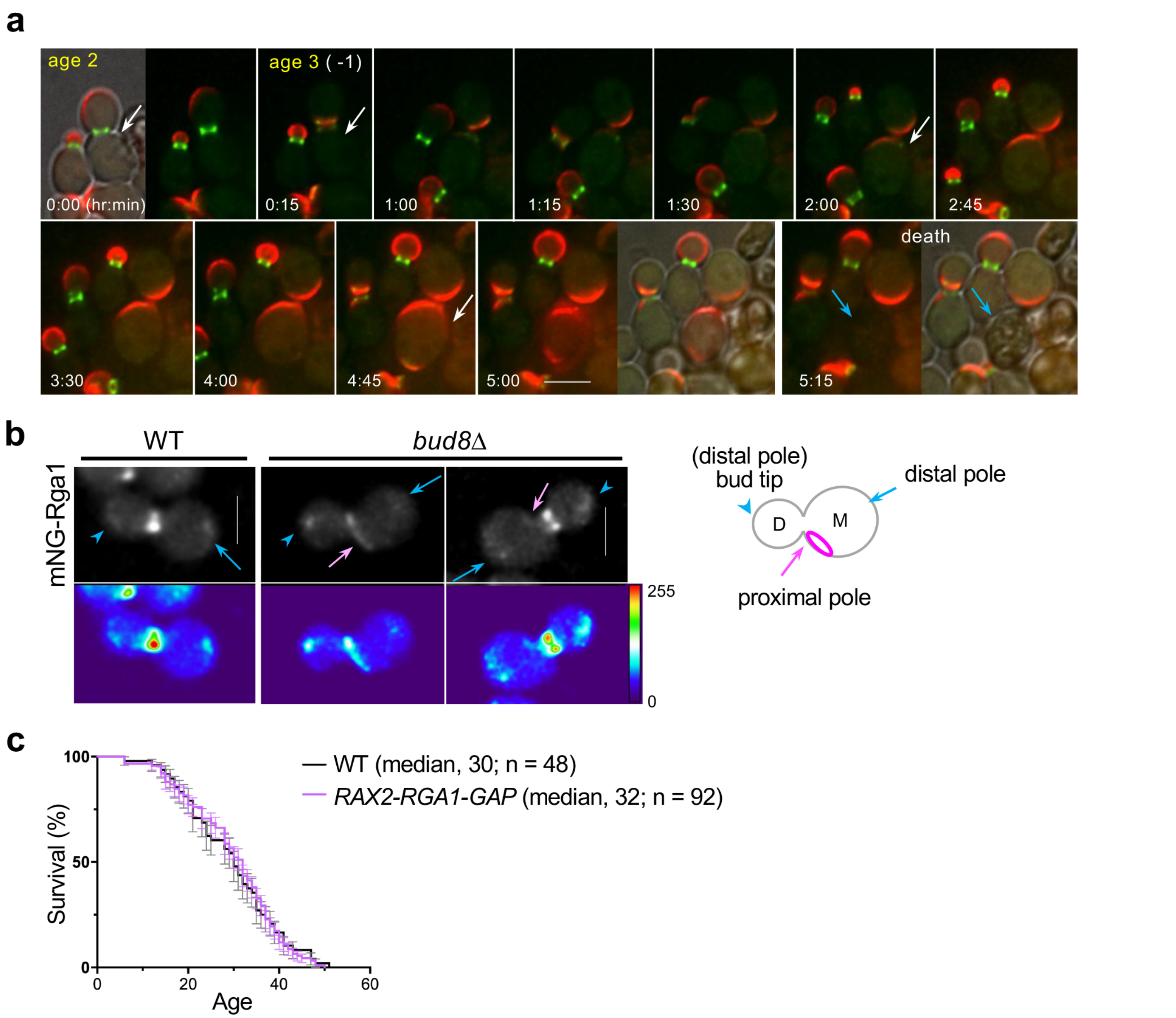


**Figure S1. Characterization of Rga1 GAP in aging**

**a.** A representative *rga1*Δ cell (expressing PBD-RFP and Cdc3-GFP) showing two Cdc42-GTP clusters before death. Ages are marked in yellow letters (-1 representing age relative to the last cell division), and selected timepoints (hr: min) are shown until death. Arrows mark the same mother cell until death (marked with blue arrows). See also Fig. 2. Scale bar, 5 µm.

**b.** Localization of mNG-Rga1 in WT and *bud8*Δ cells at young ages, shown with heatmap histograms below. Arrows and arrowheads mark the enrichment of mNG-Rga1 signals at distinct locations as depicted on the right. Scale bars, 5 µm.

**c.** The percentage (mean ± SEM) of surviving cells after the indicated number of divisions is shown for WT and a strain expressing the Rga1-GAP fused to Rax2 (see Table S1). The median survival age of each strain is indicated in parentheses: WT ($n$ = 48), *RAX2-RGA1-GAP* ($n$ = 92). Log-rank (Mantel-Cox) test indicates that the overall survival curves are not significantly different ($p$ = 0.91).

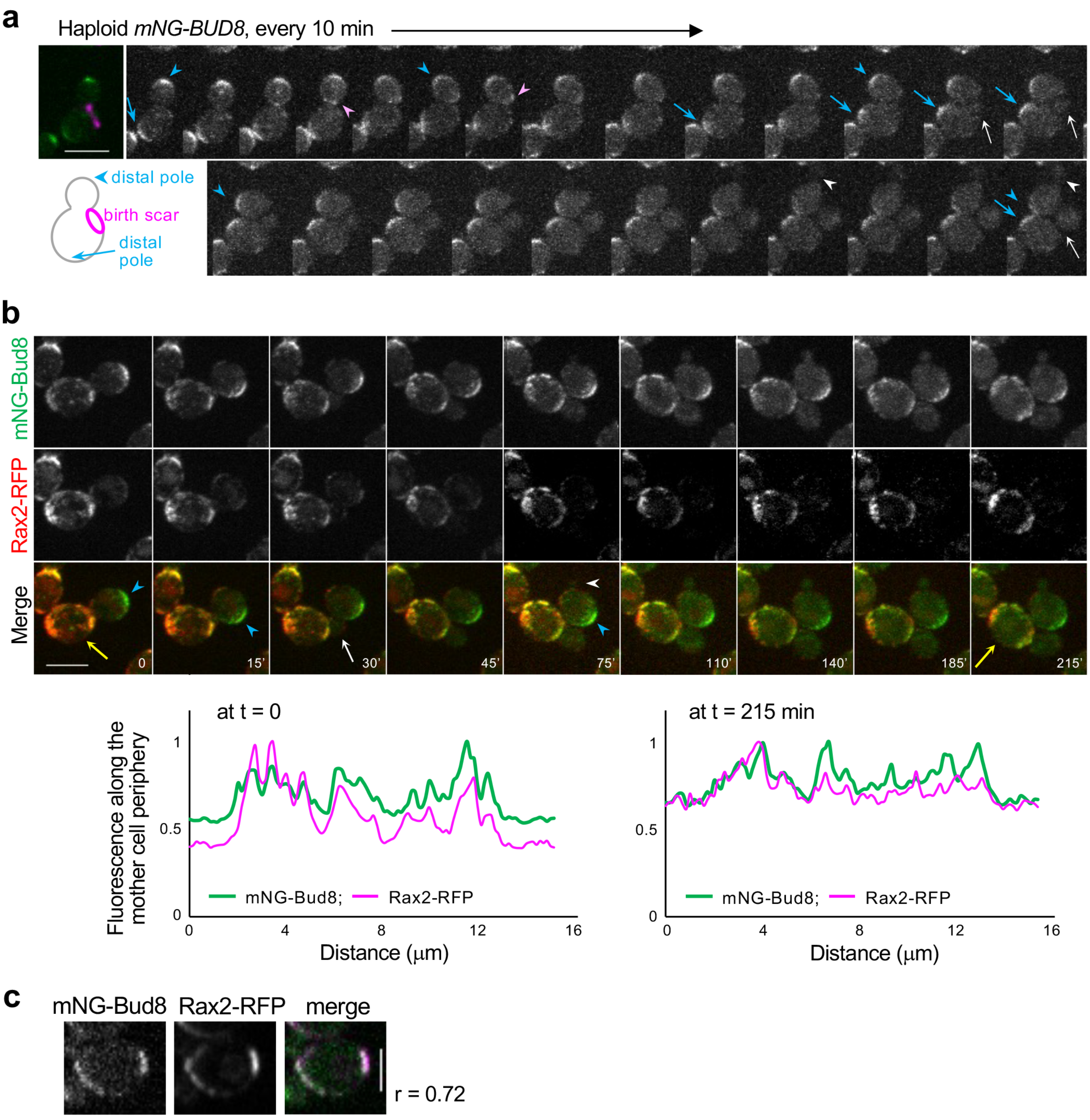


**Figure S2. Time-lapse images of mNG-Bud8 and Rax2-tdTomato**

**a.** Localization of mNG-Bud8 in a haploid WT cell, which is pre-stained with WGA-Texas Red, at 30°C. Blue arrows and arrowheads mark the distal pole of mother and bud (which became a newborn daughter cell), respectively. Pink arrowheads mark Bud8 signal at the proximal pole of the daughter cell. White arrow and arrowheads mark new buds formed from the initial mother and daughter cells, respectively. Scale bar, 5 μm.

**b.** Time-lapse images of mNG-Bud8 and Rax2-tdTomato. Blue arrowheads mark the distal pole of a bud (and a newborn daughter cell). A white arrow and a white arrowhead mark new bud emergence in mother and daughter cells, respectively. Colocalization of Bud8 and Rax2 in mother cells appears in multiple puncta at CRMs. Selected timepoints are shown. Scale bar, 5 µm. Fluorescence intensity of Bud8 and Rax2, normalized to each maximum value, is shown along the periphery of the mother cell (marked with yellow arrows) at specific time points as indicated.

**c.** Colocalization of mNG-Bud8 and Rax2-RFP in a mother cell. Pearson's correlation coefficient r (in mother cells) = 0.72. Scale bar, 5 µm.

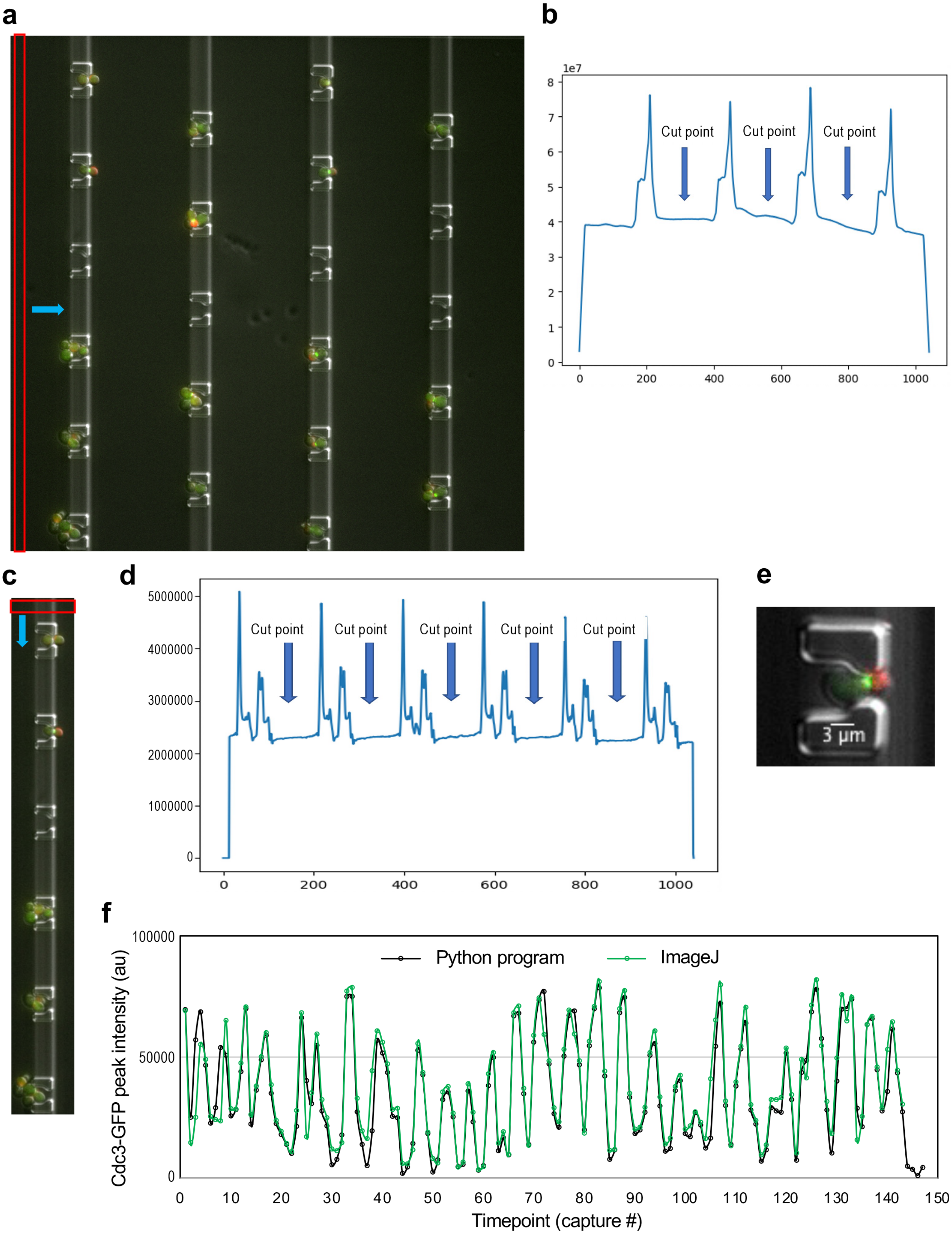
a
b
1e7
Cut point
Cut point
Cut point
c
d
Cut point
Cut point
Cut point
Cut point
Cut point
e
3 µm
f
Python program
ImageJ
Cdc3-GFP peak intensity (au)
Timepoint (capture #)

**Figure S3. A Python program and RLS estimation**

To crop the whole image (**a**), first, a window (red box) moving horizontally (blue arrow) is used. The average pixel intensity for the window is shown in (**b**). Cutting between peaks in (**b**) generates image slices shown in (**c**). Next, sliding a window (red box in **c**) vertically generates the average intensity shown in (**d**). Cutting between peaks as marked in (**d**) results in a single cell image (**e**) and the intensity values of each fluorescence channel. A cell (expressing PBD-tdTomato and Cdc3-GFP) loaded in a single trap is shown in (**e**). (**f**) The highest pixel intensity of Cdc3-GFP in a WT cell is quantified at each time point by either the Python program or ImageJ. The number of valleys of this graph represents the number of cell divisions that this cell has undergone until death.
